# Neural Oscillations during Perceptual Grouping: Insights from Alpha and Theta Bands in Visual Perception Tasks

**DOI:** 10.1101/2025.08.13.670064

**Authors:** Shefali Gupta, Tapan Kumar Gandhi

## Abstract

Perceptual grouping, the brain’s ability to organize visual stimuli into coherent objects and patterns, is thought to rely on distinct neural oscillations across various frequency bands. However, the specific contributions of these frequency bands to perceptual grouping remain a central question in cognitive neuroscience. Our study employs spectral analysis and time-frequency analysis techniques to analyze EEG data collected during visual perception tasks. By examining the temporal dynamics and frequency-specific activity, particularly focusing on the alpha and theta bands immediately following stimulus onset, our goal is to uncover the neural processes underlying perceptual grouping. Our findings indicate significant changes in alpha and theta band activity associated with perceptual grouping in the brain. These results suggest that alpha band oscillations may play a role in directing attention and filtering relevant visual information, while theta band oscillations might facilitate the integration of visual elements into coherent percepts by engaging memory processes and enhancing neural communication across brain regions. This research contributes to a deeper understanding of how neural oscillations coordinate to support perceptual organization, potentially informing future studies and therapeutic approaches aimed at enhancing visual perception and cognitive function.

## I. Introduction

Perceptual grouping is a crucial process in visual perception, enabling the brain to organize complex visual stimuli into coherent objects and patterns [1]-[6]. This capability is essential for interpreting and interacting with the environment, impacting activities ranging from basic navigation to complex tasks requiring high levels of visual discrimination [7]-[12]. The importance of frequency bands in perceptual grouping is a key area of investigation in cognitive neuroscience. Different frequency bands of neural oscillations are thought to play distinct roles in processing and integrating sensory information. However, the contribution of different frequency bands to perceptual grouping remains an open question in cognitive neuroscience. Although the alpha (8-12 Hz) and theta (4-7 Hz) bands have garnered significant attention for their potential contributions to perceptual grouping [13]-[14], their specific roles in perceptual grouping are not fully understood.

The alpha band is believed to play a role in selective attention and filtering relevant visual information, potentially aiding in the suppression of irrelevant stimuli to form coherent perceptual groups [15]. The theta band is associated with memory processes and inter-regional communication, possibly facilitating the integration of visual information across different brain regions [16]. Moreover, the precise mechanisms and interactions between these frequency bands in the context of perceptual grouping require further investigation to fully understand their contributions to this complex cognitive process [17].

Spectral analysis and time-frequency analysis are effective methods for investigating the brain’s dynamic processes. In the context of perceptual grouping, spectral analysis can reveal how different frequency bands contribute to the integration of visual information and the formation of perceptual wholes [18]. Time-frequency analysis, on the other hand, provides a dynamic view of neural activity by tracking how oscillatory power in various frequency bands changes over time, which is especially useful for examining transient neural responses to stimuli [18]-[20]. Another approach to gaining insights into the brain’s dynamic process of perceptual grouping is by tracking changes in neural activity following stimulus onset. When visual stimuli are presented, the brain quickly processes and organizes this information into coherent perceptual units. Observing these neural activity changes across different frequency bands reveals how the brain integrates and organizes visual information [21]-[25].

In summary, this paper presents an investigation into the spectral and time-frequency characteristics of neural activity associated with perceptual grouping. By leveraging the power of spectral analysis and focusing on alpha and theta band activity immediately following stimulus onset, we aim to uncover the oscillatory dynamics that enable the brain to create coherent perceptual experiences from fragmented visual inputs. This work contributes to the broader field of cognitive neuroscience by providing new insights into the frequency-based mechanisms of visual perception.

## II. Materials and Methods

### A. Participants

Fifteen subjects aged 19 to 30 participated in the study, including 13 males and 2 females. None of the subjects had any neurological deficits or illnesses. Participants were fully informed about the experiment, and consent was obtained prior to participation. The experiment was approved by the Indian Institute of Technology Ethical Clearance Committee.

### B. Experiment Protocol

EEG signals related to visual stimuli were recorded using the ActiChamp Brain Vision device with a sampling frequency of 2500 Hz. A 64-channel EEG acquisition system captured the raw data, employing electrodes placed according to the international 10-20 system and headcap size tailored to participants’ head circumferences. Signals were transmitted to the Brain Vision amplifier for processing. Participants sat 40-60 cm from a monitor, maintaining low scalp-electrode impedance. They were instructed to minimize movement and focus on the screen during the experiment duration.

### C. Experiment Design

The experiment featured two types of images: ‘Structure’ and ‘Non-Structure’. In the ‘Structure’ images, random dots were arranged to create the appearance of two straight lines, whereas the ‘Non-Structure’ images consisted solely of random dots without forming any recognizable patterns. Each image appeared on the screen for 1000 ms, separated by a random interval ranging from 1200 ms to 2300 ms. A total of 40 images were used, with each stimulus presented randomly five times, resulting in a total of 200 events (100 ‘Structure’ and 100 ‘Non-Structure’). Fig. 1 depicts the experiment design outlook.

**Fig. 1:**
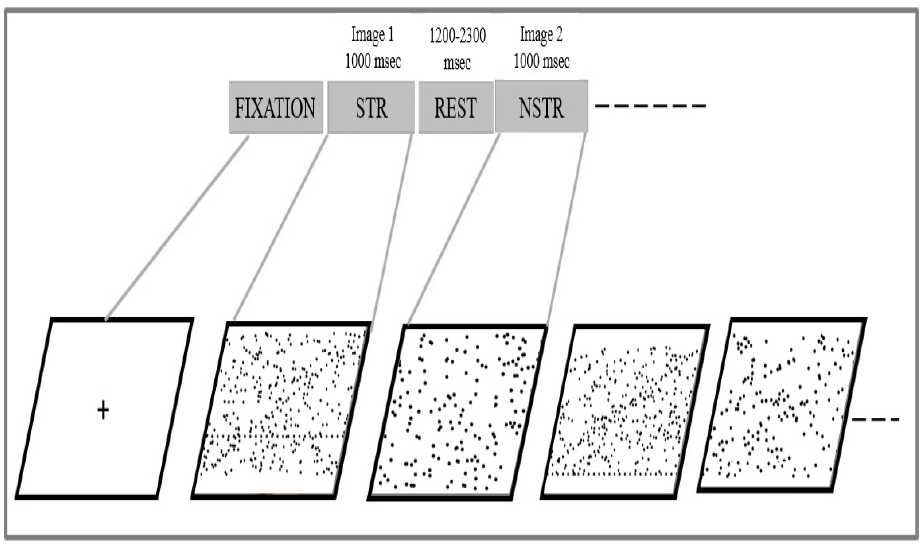
Outlook of stimulus presentation: Each image was displayed on the screen for 1000 ms, followed by an interstimulus interval ranging from 1200 ms to 2300 ms between successive images. In total, 200 stimuli, comprising 100 structure images and 100 non-structure images, were presented randomly to the participants. The image displays random dots, with some appearing to create perceptual line segments referred to as ‘Structures’ (STR) while the other image solely consists of random dots, referred to as ‘Non-Structures’ (NSTR).

### D. Analysis

#### 1) Preprocessing

The raw EEG data underwent preprocessing steps that included bandpass filtering with cutoff frequencies of 2 Hz to 45 Hz. Each signal was then re-referenced using average re-referencing before epoch extraction. Epochs were segmented to capture events related to both structure and non-structure stimuli, using a window length of 1400 ms from −200 ms before stimulus onset to 1200 ms after stimulus onset. The mean baseline recorded from −200 ms to 0 ms was subtracted from each channel to remove baseline effects.

#### 2) Time-Frequency Analysis

Time-Frequency analysis using Continuous Wavelet Transform (CWT) allows for timevarying frequency analysis, to examine how the frequency content of a signal changes over time. This is particularly useful in EEG studies, where neural responses to stimuli vary over time. By revealing when and how different frequency components are activated in response to visual stimuli, CWT enhances our understanding of how the brain processes and integrates information during perceptual tasks.

The time-frequency map was computed for the two events—structure and non-structure separately by averaging over all electrodes of interest. The significance between events was checked using ANOVA.

**Fig. 2:**
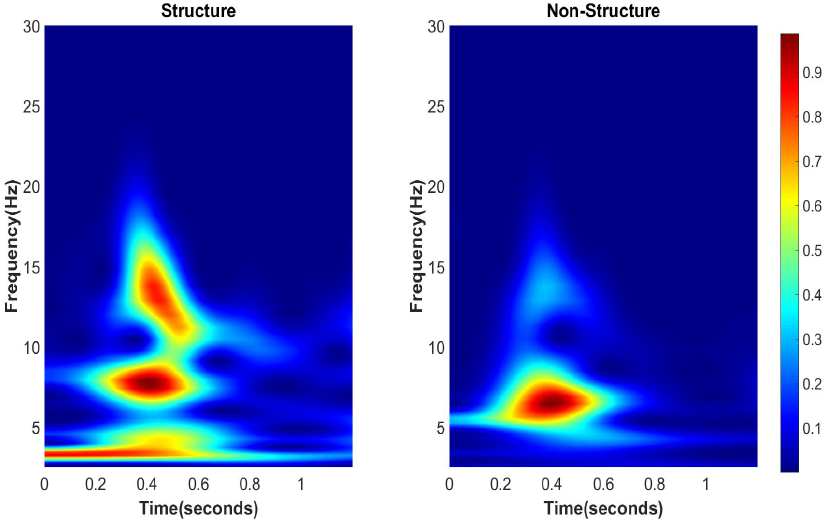
Time Frequency Analysis for the events-Structure and Non-Structure. The result reflects more activation for perceptual grouping in the Theta and Alpha bands.

#### 3) Spectral Analysis

Spectral analysis is a process used to examine the frequency content of time-series signals, such as those recorded in EEG or MEG studies. It allows researchers to break down complex signals into their constituent frequencies and analyze how these frequencies vary over time or across different conditions. Fast Fourier Transform (FFT) algorithm was employed to convert the time-domain signal into the frequency domain.

#### 4) Power Analysis

Power was calculated at alpha and theta bands for the two classes of task. The significance of the difference between the classes was checked using ANOVA. Calculating changes in power before and after stimulus onset is a common approach in spectral analysis to understand how brain activity is modulated by stimuli. Power pre-stim and post-stim onset were calculated for both events at theta and alpha bands. The significance was measured using ANOVA.

**Fig. 3:**
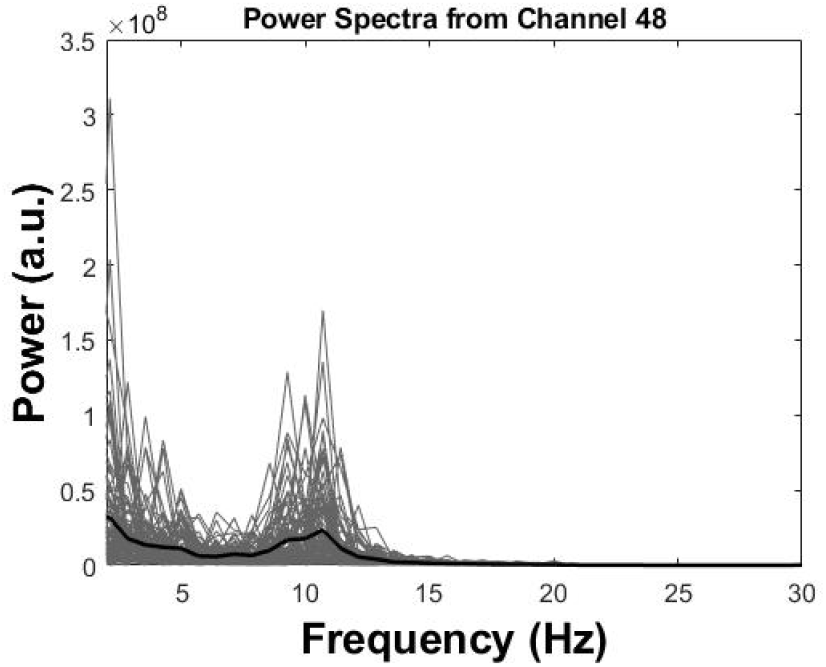
Spectral analysis of EEG data showing power distribution across frequency bands. The Spectra highlight changes in neural oscillatory activity, with notable increases in power within the theta and alpha band, indicating a cognitive process of perceptual grouping.

## III. Results

### A. More activation for perceptual grouping in theta and alpha band

The spectral analysis performed for all structure epochs indicates more activation in theta and alpha band for perceptual grouping. The Time-Frequency Analysis displays enhanced connectivity in theta and alpha bands for structures. Enhanced alpha activity suggests heightened attention and inhibition of irrelevant information, facilitating focused perceptual processing. Concurrently, increased theta activity points to the brain’s engagement in integrating and organizing sensory inputs, reflecting cognitive processes essential for forming coherent perceptual experiences.

### B. Post-Stimulus Theta Band Activity Change for structures in Frontal Brain region

The alteration in post-stimulus activity within the frontal brain region, particularly in the theta frequency band was observed (as indicated by topographs). Theta oscillations in the frontal cortex are linked to the brain’s initial processing of sensory information, sustaining cognitive processes, and facilitating communication between different brain regions.

### C. Post-Stimulus Alpha Band Activity Change in the occipital region for structures

After the stimulus onset, the alterations in alpha band activity were observed particularly in the occipital brain region (as indicated by topographs). Correlation Analysis between Power at 10 hz and task-related alpha power evaluated reflected high value of correlation coefficient ensure the involvement of alpha activity in perceptual grouping task.

This increase in alpha power, known as alpha synchronization, indicates a state of cortical inhibition and reduced excitability, suggesting that the brain is filtering out irrelevant information to focus on the pertinent visual input. This phenomenon helps in enhancing selective attention and visual perception by suppressing background noise and optimizing the processing of the stimulus.

## IV. Discussion

Perceptual grouping, the process by which the brain organizes visual stimuli into coherent objects and patterns, is facilitated by distinct neural oscillations across various frequency bands. Alpha (8-12 Hz) and theta (4-7 Hz) bands play significant roles in this process. The alpha band is associated with attention and controlled access to stored information, helping to filter relevant visual information and suppress irrelevant inputs. This selective attention mechanism is crucial for forming coherent perceptual groups. The theta band, on the other hand, is linked to memory processes and inter-regional communication within the brain. Theta oscillations modulate higher-frequency gamma band activity, which is essential for synchronizing different brain regions during perceptual tasks. This cross-frequency coupling ensures efficient integration of visual information, enabling the brain to construct unified perceptual experiences from fragmented sensory inputs. Thus, the interplay between alpha and theta bands, along with their interactions with other frequency bands, underpins the neural dynamics of perceptual grouping.

The findings from our study provide compelling insights into the role of neural oscillations, particularly in the alpha and theta bands, during perceptual grouping in visual processing tasks. Our use of spectral and time-frequency analysis techniques on EEG data revealed significant changes in alpha and theta band activity following stimulus onset, highlighting their involvement in the brain’s organization of visual stimuli into coherent percepts.

The observed increase in alpha band activity suggests a role in directing attention and filtering relevant visual information [26]-[28][15]. This aligns with previous research indicating that alpha oscillations are involved in suppressing irrelevant sensory inputs, thereby enhancing the processing of pertinent stimuli. In the context of perceptual grouping, heightened alpha activity post-stimulus onset may reflect the brain’s selective engagement with visual features crucial for forming cohesive perceptual units.

Furthermore, our findings indicate dynamic changes in theta band oscillations, implicating their role in memory processes and neural communication across brain regions [29]-[31][16]. Theta band activity may facilitate the integration of visual elements by coordinating the retrieval and comparison of stored representations, which is essential for assembling coherent perceptual groups. This supports the notion that theta oscillations contribute to the temporal binding of visual information, enabling the brain to construct meaningful interpretations of sensory inputs.

**Fig. 4:**
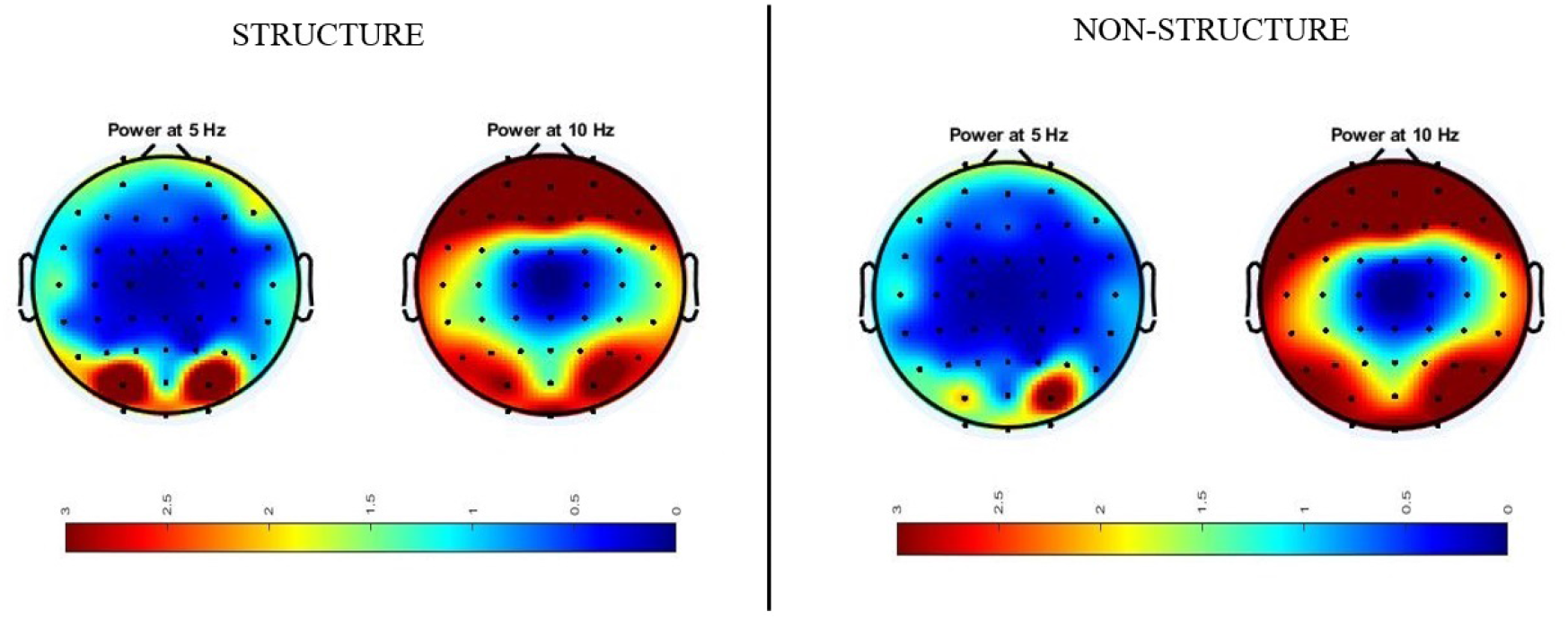
Topographs displaying the Power at Theta and Alpha band for the events-Structure and Non-Structure

**Fig. 5:**
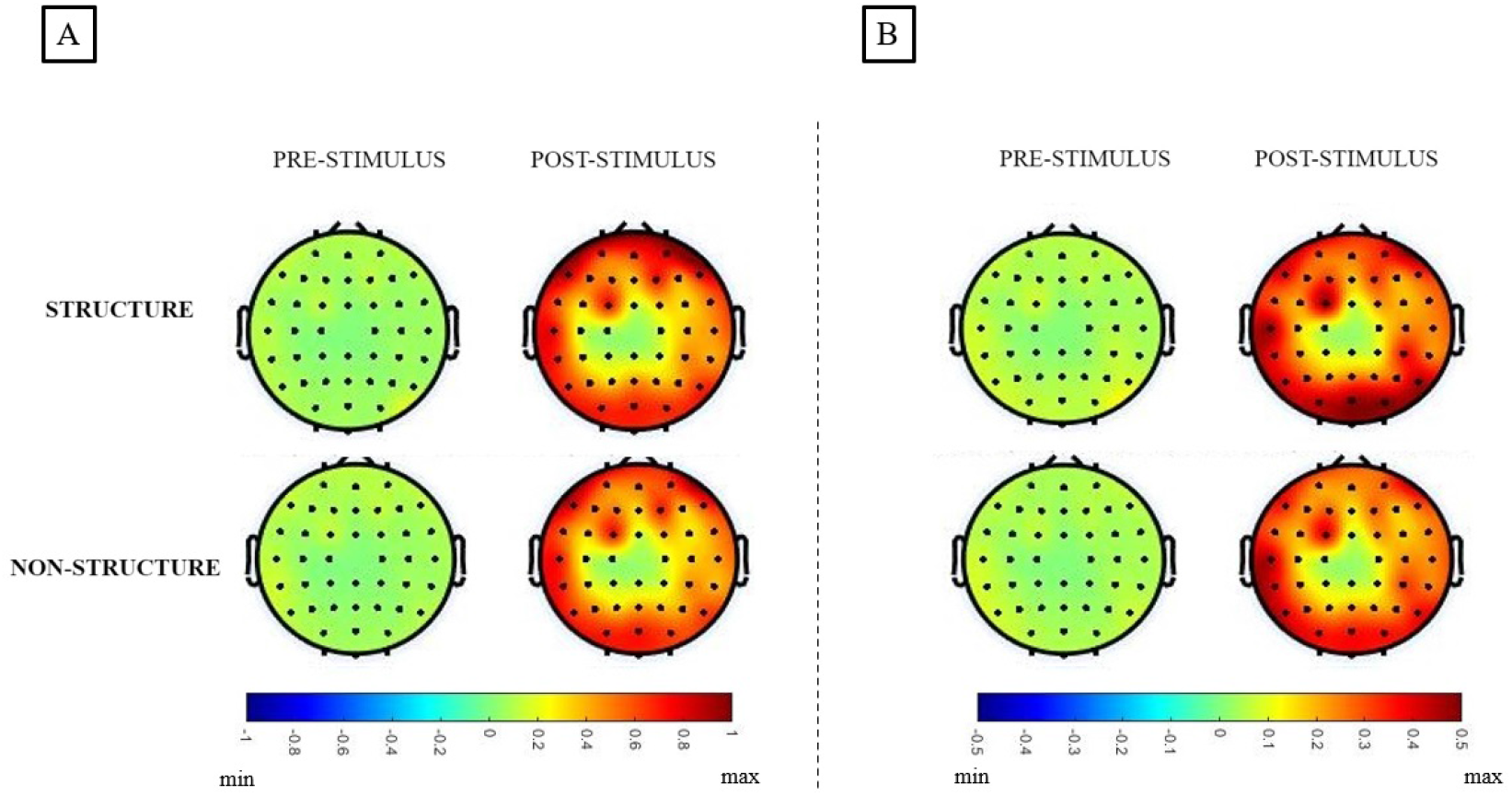
(A) Topography maps representing the Theta Power pre-stimulus (before stimulus onset) and post-stimulus (just after stimulus onset) (B) Topography maps representing the Alpha Power pre-stimulus (before stimulus onset) and post-stimulus (just after stimulus onset).

The integration of alpha and theta band dynamics observed in our study underscores their complementary roles in perceptual organization. Alpha oscillations may initiate selective attention mechanisms, while theta oscillations coordinate memory-related processes that aid in integrating and interpreting visual stimuli [32]-[34][17]. These findings provide a nuanced understanding of how neural oscillations coordinate within the brain to support perceptual grouping, offering potential implications for enhancing both visual perception and cognitive function through targeted interventions.

Moving forward, future research could explore the specific mechanisms through which alpha and theta band activities interact during different stages of perceptual processing. Additionally, investigating how these neural oscillations are modulated under varying experimental conditions or in clinical populations could further elucidate their contributions to perceptual organization and inform therapeutic strategies aimed at improving visual and cognitive abilities.

## V. Conclusions

Our study highlights the critical roles of alpha and theta band oscillations in perceptual grouping, as evidenced by changes in EEG activity following visual stimulus onset. Alpha band activity appears crucial for selective attention and enhancing relevant sensory processing, while theta band oscillations facilitate memory retrieval and integration processes essential for assembling coherent perceptual units. These findings deepen our understanding of how neural oscillations coordinate to support visual perception, offering insights into potential avenues for enhancing cognitive function and visual processing through targeted interventions. Further research exploring the interactions between alpha and theta band activities in various contexts could provide additional clarity on their specific contributions to perceptual organization and inform therapeutic approaches for improving cognitive and visual outcomes in clinical settings.

**Fig. 6:**
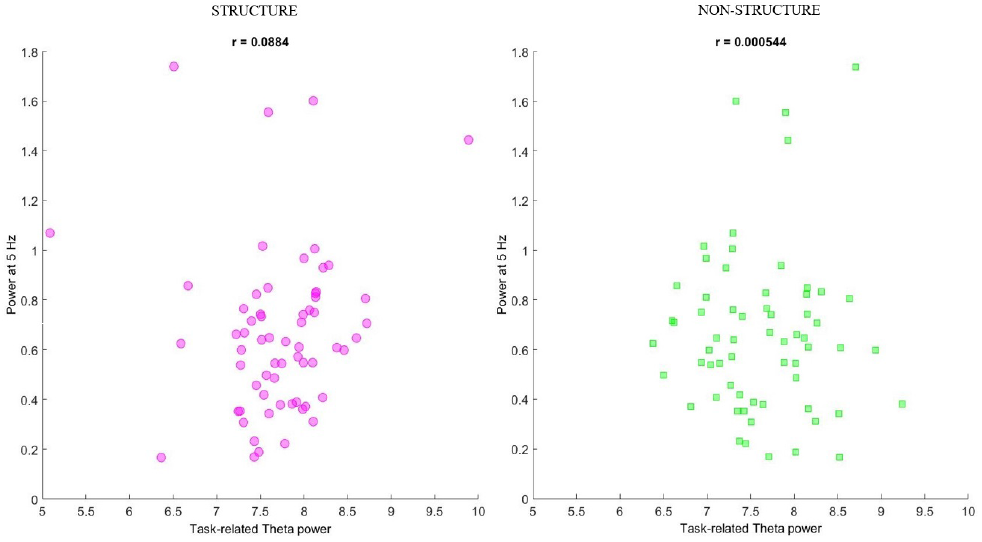
Correlation between Power at 5 Hz and Task-related theta power for the events Structure and Non-Structure.

**Fig. 7:**
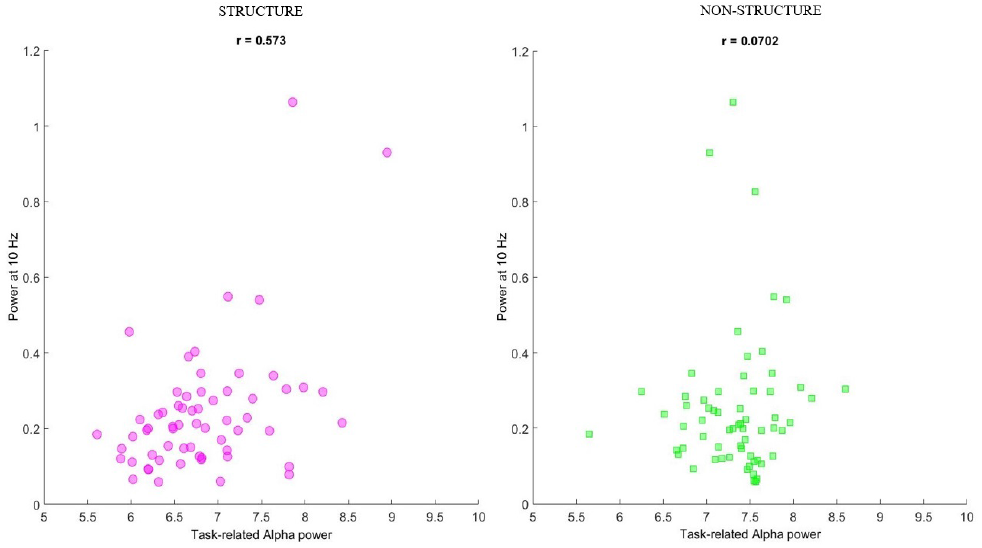
Correlation between Power at 10 Hz and Task-related alpha power for the events Structure and Non-Structure.

## CRediT authorship contribution statement

Shefali Gupta: Conceptualization, Data Curation, Methodology, Analysis, Writing and Editing– original draft. Tapan Kumar Gandhi: Methodology, Supervision, Editing – original draft.

## Acknowledgements

The authors extend their gratitude to all participants who contributed to the study for their cooperation, which facilitated smooth data collection.

## Data and code availability

The data that supports the findings of this study is available from the corresponding author upon reasonable request. The code will be made available on request.

